# Quantitative monitoring of nucleotide sequence data from genetic resources in context of their citation in the scientific literature

**DOI:** 10.1101/2021.05.10.443393

**Authors:** Matthias Lange, Blaise T.F. Alako, Guy Cochrane, Mehmood Ghaffar, Martin Mascher, Pia-Katharina Habekost, Upneet Hillebrand, Uwe Scholz, Florian Zunder, Jens Freitag, Amber Hartman Scholz

## Abstract

**Background:** Linking nucleotide sequence data (NSD) to scientific publication citations can enhance understanding of NSDs provenance, scientific use, and re-use in the community. By connecting publications with NSD records, NSD geographical provenance information, and author geographical information, it becomes possible to assess the contribution of NSD to infer trends in scientific knowledge gain at the global level.

**Findings:** For this data note, we extracted and linked records from the European Nucleotide Archive to citations in open-access publications aggregated at Europe PubMed Central. A total of 8,464,292 ENA accessions with geographical provenance information were associated with publications. We conducted a data quality review to uncover potential issues in publication citation information extraction and author affiliation tagging and developed and implemented best-practice recommendations for citation extraction. Flat data tables and an data warehouse with an interactive web application were constructed to enable ad hoc exploration of NSD use and summary statistics.

**Conclusions:** The extraction and linking of NSD with associated publication citations enables transparency. The quality review contributes to enhanced text mining methods for identifier extraction and use. Furthermore, the global provision and use of NSD enables scientists around the world to join literature and sequence databases in a multidimensional fashion. As a concrete use case, statistics of country clusters were visualized with respect to NSD access in the context of discussions around digital sequence information under the United Nations Convention on Biological Diversity.

## Data Description

Nucleotide Sequence Data (NSD) plays a fundamental role in biological research ranging from public health and medical applications to understanding the molecular basis of life and evolution, such as how genes (mis)function in disease mechanisms [1], insights into ecosystem functioning and biodiversity conservation, and to assist in breeding new plant variety and animal breeds enabling food security and sustainability [2]. Scientifically, NSD plays a major role for mechanistic modeling of species evolution [3], genotype – phenotype correlation [4], to identify and mitigate risks to species, track their illegal trade, identify the geographical origin of products, and plan conservation management strategies [5].

These applications demonstrate the wide value of NSD use and application and have triggered political debate about benefit sharing from genetic resources (GR). Under the Convention on Biological Diversity (CBD) and the Nagoya Protocol [6] as well as the International Treaty on Plant Genetic Resources for Food and Agriculture (ITPGRFA) the topic of “digital sequence information” (DSI) has garnered immense interest and raised concern across the international scientific community. Discussions have focused on the use of NSD from GRs, as DSI is an undefined and non-scientific term. Due to the exponential growth of public sequence and downstream databases [7] many parties are concerned that insufficient benefit-sharing is taking place. Datasets such as this one provides an opportunity for evidence-based policymaking to analyze global trends in NSD provision and use as well as other science policy fields, including scientific strategic development and internationalization.

### Context

FAIR (findable, accessible, interoperable, re-usable) data principles as defined in 2016 in the FAIR Guiding Principles for scientific data management and stewardship [8] guide the design of open data sharing infrastructures as an enabling technology for economic and scientific progress. Data sharing principles have been implemented at national and international levels. For example, The German Federal Ministry of Education and Research (BMBF) has funded an interdisciplinary project called “Science-based approaches for Digital Sequence Information” (WiLDSI) [9] which aims to (i) raise awareness and involve the international scientific community into the debate and decision making process surrounding DSI, (ii) to identify and elaborate scenarios for open access to the NSD and (iii) to establish fair and sustainable benefit sharing.

In this context, transparent quantitative measures of NSD citation and re-use can inform decision-making processes surrounding the design of data sharing infrastructure, awarding scientific “credit” or political acknowledgment, or addressing needs of commercial users [10]. Data citation has received increased attention from publishers, funding agencies and infrastructure providers [11] [12] in recent years. However, best practices for NSD citation are still lacking and those developed for scientific publications cannot be readily transferred. This is true especially for NSD, which is hosted by the core data infrastructure, the International Nucleotide Sequence Database Collaboration (INSDC) [13]. The European Nucleotide Archive (ENA) [14] and Europe PubMed Central (ePMC) [15] are, respectively, the European partners in INSDC and a repository of open access articles. Both have a long tradition of handling open data and document the heterogeneous quality of author’s data citation practices [16]. ePMC listed publications generally employ text-embedded ENA identifiers, like accession numbers, project accessions, or study accessions.

## Methods

Figure 1 shows the extraction of ENA citations performed in three phases. First, ENA accessions and project accession numbers were extracted. Literature citations listed directly in the ENA entry were extracted in parallel and are called herd “primary publications”. Next, we retrieved scientific papers that referred to these accession IDs via a full text search using the ePMC REST API [17]. These publications we labelled “secondary publications”. Finally, the extracted references and associated citation information were organized into six tables (Figure 2) and imported into a data warehouse.

**Figure 1.**
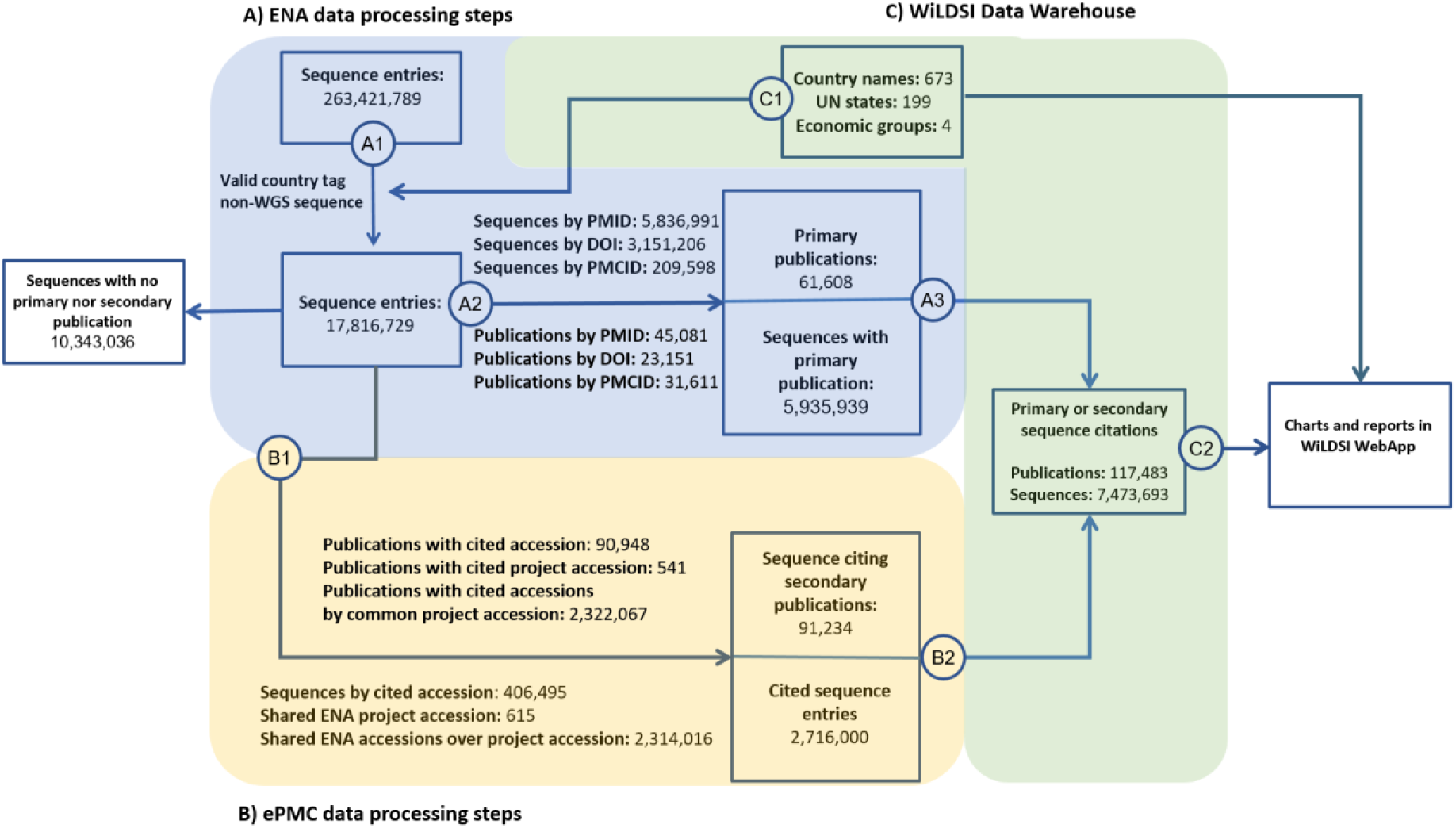
Visualization of data processing performed to extract, filter and join ENA and ePMC data sets. ENA records are parsed (A1) and filtered for valid country tag and fed into ePMC RestFul API to extract matching secondary publication (B1) by ENA accession or project accession numbers. Primary publications are linked by ENA record (A2) to the DOI, PMCID or PMID. The resulting data sets are normalized as tables ENA_SEQUENCES, PMC_REFERENCES and loaded into the data warehouse (A3, B2) alongside a curated list of world’s countries in table COUNTRIES and economics groups in table COUNTRY2GRP (C1). SQL queries (C2) are applied to generate charts and reports in the Web application.

**Figure 2.**
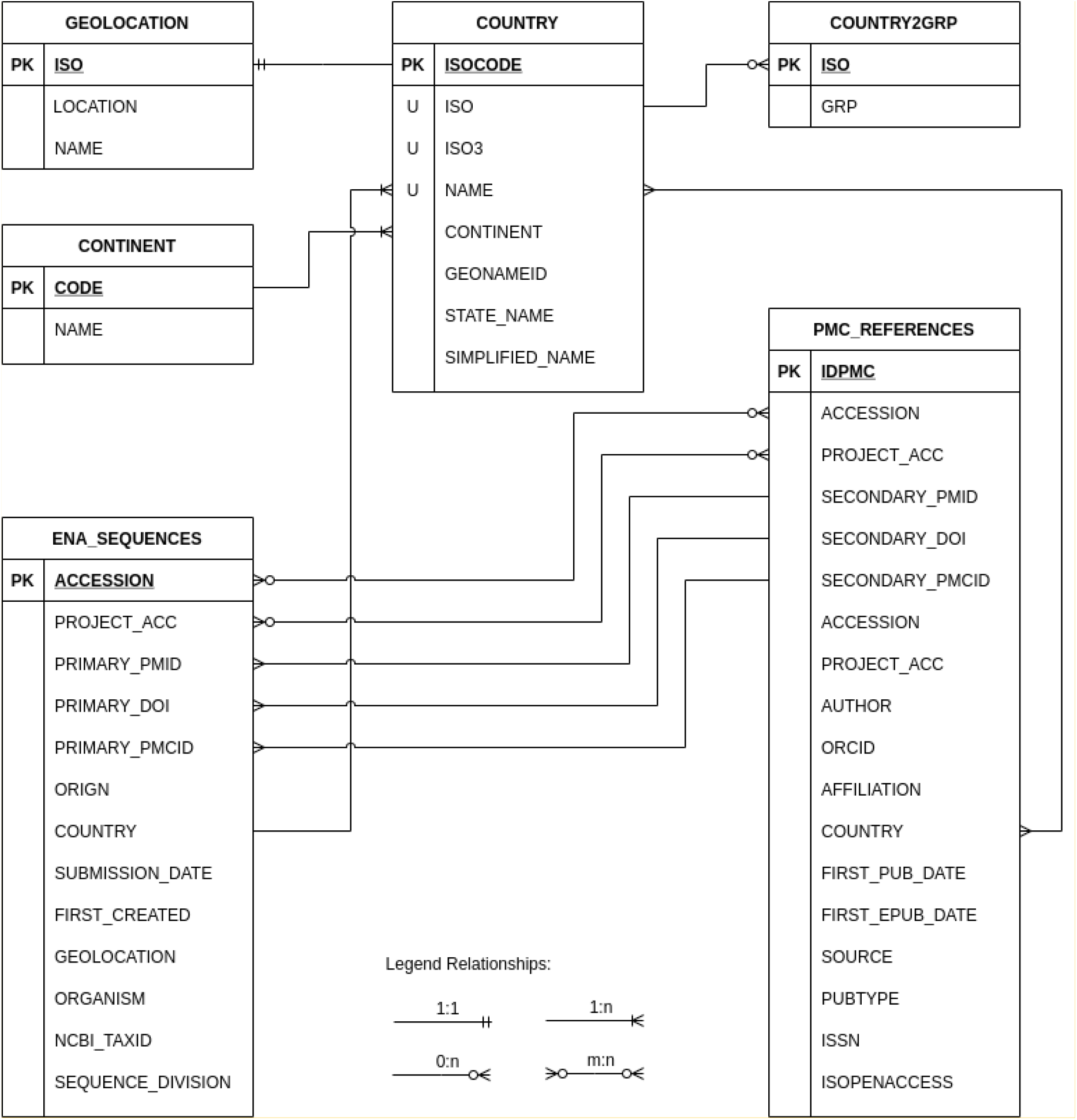
Table schema of the WiLDSI data warehouse - The table ENA_SEQUENCES comprise metadata of a sequence stored in EBI ENA database. The attributes accession and project accession are used to join secondary literature that cite sequences. The attribute country refers to the country table to resolve and group country tagged ENA sequences. The table PMC_REFERENCES consists of all ePMC published papers either referencing a ENA sequence by accession or project accession and references from ENA records as primary publication by either a DOI, PMID or PMCID.

Five classes of citation pattern are used for ePMC publication as ENA identifiers where used: single accession number using word separation characters, e.g. hyphens, brackets, quotation marks; range notation of referenced accessions, text embedded enumeration, lists in supplement material or even embedded into figure bitmaps. The data extraction process from EMBL-ENA and ePMC was executed by Perl and Python scripts. The starting point of the data extraction process is the EMBL-ENA flat file dump of release 143, which was obtained from the EMB-ENA FTP server and comprised 263,421,789 records. Next, all non-WGS ENA records were parsed to compile a relevant set of attributes for the table “ENA_SEQUENCES”. A total of 17,820,136 ENA accessions with valid country tags (i.e., the /country field in the ENA entry, comprising 15% of all records) were included. Next, using the ePMC REST API, these ENA accessions were scanned in 36.7 million full text articles accessible via PubMed. Due to performance reasons, this text tokenization was executed on site at EBI in a compute cluster environment. From the resultant publications those were select that have valid author country information and that either (b) cite an ENA sequence as a secondary publication^1^ or (b) that are cited by ENA record as primary publication^2^. The publications matching these criteria were compiled into the table “PMC_REFERENCES”. In detail, 5,935,939 sequences cite 61,608 publications and 2,716,000 sequences are cited by 91,234 publications respectively. All scripts used in our analyses are available in GitHub [18].

The table “COUNTRY” was compiled and curated from UN state membership [19]. It comprises the three kinds of ISO-3166-1-codes, the official name (e.g. United Kingdom of Great Britain and Northern Ireland), a short version of the state name (e.g United Kingdom), commonly used names (e.g. Great Britain), and continent assignments. This table allows mapping from partly ambiguous country affiliation used in papers to the actual country designations recognized under international law. In particular, provinces or (partially) autonomous areas, such as Taiwan or West Sahara, are mapped to the legally responsible UN state party. Furthermore, several Ocean areas, for example “Bismarck Sea” or “East China Sea” are grouped together under the “Ocean” label along with more standard fields such as “Atlantic Ocean”^3^. The assignment to economic groups is stored in table “COUNTRY2GRP”. Here a 2-letter ISO code is assigned to rough economic groups OECD (Organization for Economic Cooperation and Development), BRICS (Brazil, Russia, India, China, South Africa), and G77 (representative of developing economies). In order to visualize countries in a world map, we used the table “GEOLOCATION” comprising the coordinates of the centroid of each country.

The tables are provided for download as CSV files (see section “Data Availability”). For further processing, we imported an ORACLE SQL data warehouse that employs state-of-the-art database technologies such as in-memory tables, vectorization and columnar storage for the fast execution of complex queries on large datasets.

### Data validation and quality control

In order to assess the reliability of the extracted ePMC to ENA references, potential quality issues were evaluated by plausibility scans across data warehouse tables, including inspection of 20 randomly sampled papers performed by domain experts from IPK’s sequence submission service team. We also took into account review articles on the use of data identifiers in life science literature [20] [21]. Finally, we applied the Dimensions text mining tool [22] to cross-check the sensitivity of ePMC API in respect of recall and sensitivity, e.g., to find false negative hits such as published articles that reference ENA sequences but which were not found by the ePMC REST API.

#### Country names

Country name had to match records in the country table. Here we found some obsolete or ambiguous country names, like Montenegro or West Sahara and historic country names, like Soviet Union, which cannot be assigned uniquely to current UN states. Ambiguous country names were resolved manually and reverted to synonyms in the country table (e.g. Cote d’Ivoire to Ivory Coast amongst others). ENA or PMC records with obsolete country tags were kept in the data set but ignored for summary statistics queries and excluded in below quality check.

#### Retrieval of referenced IDs

Next, reference integrity between the tables was checked. SQL queries were applied to count unique paper identifiers, ENA accessions, and country tags. We checked whether the primary papers referenced in an ENA record exist in the ePMC databases. A total of 6,753,891 ENA records refer either by DOI, PMID, or PMCID to 351,119 ePMC records. Conversely, there are 9,589,900 ENA records without any primary literature reference. Furthermore, we confirmed that ePMC records, which cite secondary ENA accession or project numbers can be resolved to records in ENA. We found 189,581 ePMC records that reference 2,801,072 ENA records by either accession or project accession number. A potential issue in the context of using an identifier to cite ENA records is that authors sometimes use ENA study identifiers or even BioSample IDs. However, our pipeline considers ENA accession and project accession only.

#### Author identification

The combination of first and last names are not unique identifiers for human beings. ORCIDs provide unique identifiers for authors and are on their way to becoming compulsory for publications. Existing articles, however, are only occasionally associated with ORCID. Another potential issue is that it is possible to register multiple ORCID for one person. Identifying authors as a concatenation of author name and affiliations is error prone [23]. Therefore, author information was retained in the tables but not used for statistical analysis.

#### Range notation

Scientific publication may use ambiguous range notation to cite ENA accessions. As illustrated in Figure 3, hyphen as range notation aggregate a sequence of ENA accessions. Here, the authors assume an ordered sequence of accession numbers and it is interpreted as such by human readers, but is not recognized by programmatic text mining. Thus in the data extraction used here, a potentially high number of ENA accessions are missed and the dataset is an underestimate of the number of referenced DSI. This analysis is intended to support future work to address these shortcomings.

**Figure 3:**
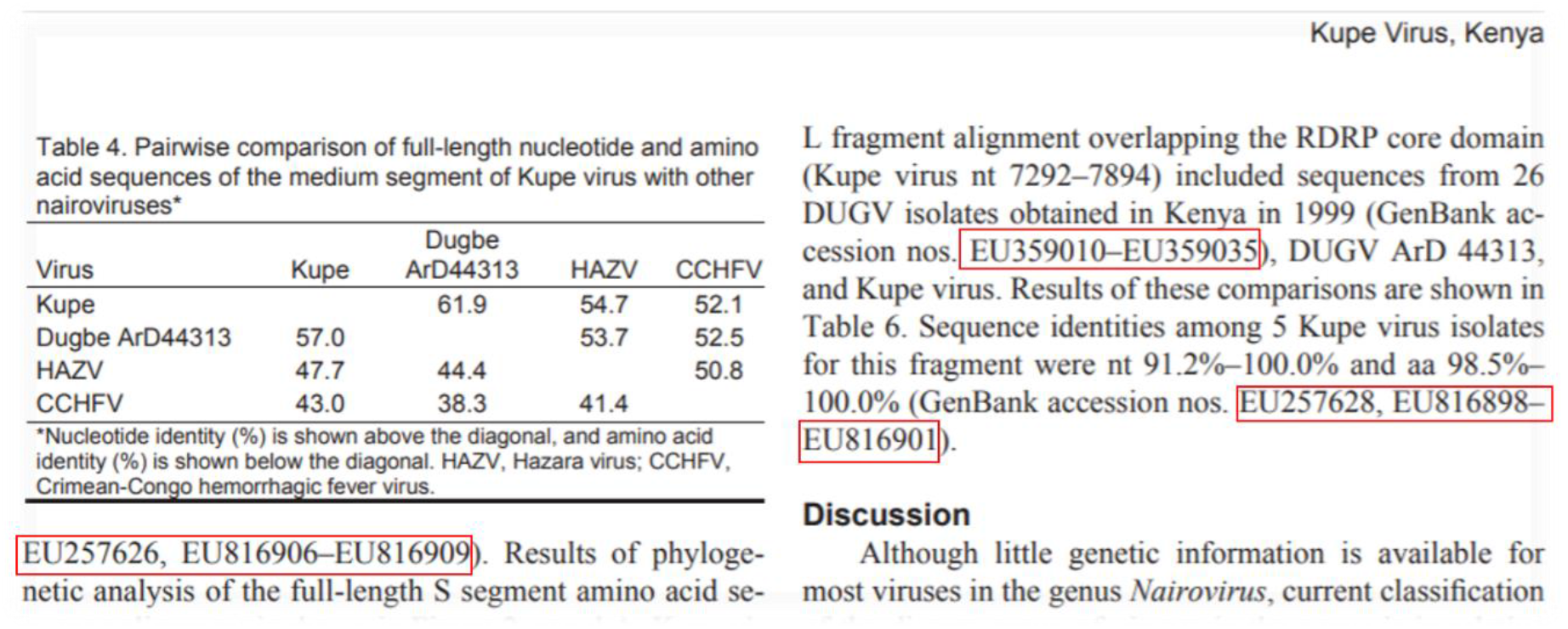
Example of range notation for ENA accession references. Within the selected part of publication with DOI: 10.3201/eid1502.080851 the actual number of cited ENA accessions is 35, but ePMC API matched 8 only.

#### Access restrictions

Only open access publications were available for high-throughput text mining. In order to efficiently process 18 million ENA accession numbers the ePMC REST API at EBI’s local compute infrastructure was used. This causes a potential loss of recall in comparison to a broad and integrative use of further state-of-the-art literature mining services that include articles behind paywalls. In order to get an estimation of potential missed DSI citation, we used alternative tools that cover patent and closed access publications. We applied the commercial “Dimensions” [22] and the free “Lens.org” [24] search tools, which include patent and restricted-access publication, to compare recall performance for 20 randomly selected ENA accessions. To work with a comparable corpus, this evaluation was performed within 4 weeks of the ePMC based text mining run. The results are compiled in Table 1. Specific hits to one of three approaches, ePMC, Lens, dimensions were observed. This is likely due to the larger corpora of Dimensions and Lens. For example, ENA accession AB076935 was linked to 3 public and 3 closed access publications by Dimensions, whereas ePMC did not report any matching publication. Differences in file format may explain some of the differences. There are cases where the PDF rendered articles differ from ePMC rendered HTML versions, so that the PDF versions can contain more ENA accession numbers than HTML versions in ePMC. We did not aim for an in-depth analysis for literature search tools, but our cursory overview supports the notion that a substantial number of publications relevant to NSD may be behind paywalls. However, as the spirit of the project in which this analysis took place, with a heavy emphasis on open access, we continued our analyses with the open dataset.

**Table 1.**
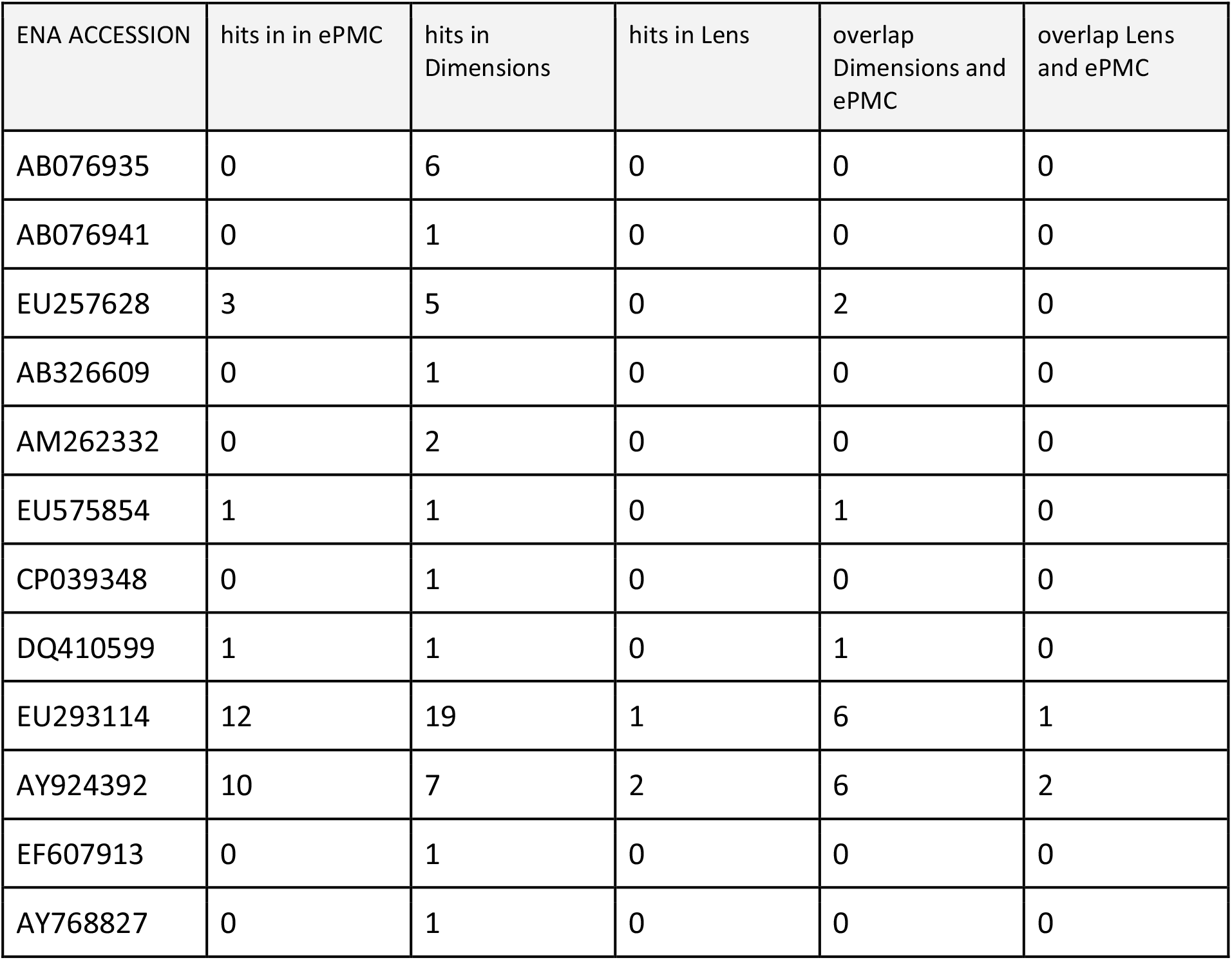
Comparison of ENA accession number query APIs performance of APIs of EBI ePMC, Dimensions^4^ and Lens^5^.

| ENA ACCESSION | hits in in ePMC | hits in Dimensions | hits in Lens | overlap Dimensions and ePMC | overlap Lens and ePMC |
| --- | --- | --- | --- | --- | --- |
| AB076935 | 0 | 6 | 0 | 0 | 0 |
| AB076941 | 0 | 1 | 0 | 0 | 0 |
| EU257628 | 3 | 5 | 0 | 2 | 0 |
| AB326609 | 0 | 1 | 0 | 0 | 0 |
| AM262332 | 0 | 2 | 0 | 0 | 0 |
| EU575854 | 1 | 1 | 0 | 1 | 0 |
| CP039348 | 0 | 1 | 0 | 0 | 0 |
| DQ410599 | 1 | 1 | 0 | 1 | 0 |
| EU293114 | 12 | 19 | 1 | 6 | 1 |
| AY924392 | 10 | 7 | 2 | 6 | 2 |
| EF607913 | 0 | 1 | 0 | 0 | 0 |
| AY768827 | 0 | 1 | 0 | 0 | 0 |

## Re-use potential

To enable the further exploration of the data set, a web application was developed and is publicly accessible at https://wildsi.ipk-gatersleben.de. We focused especially on understanding NSD/DSI usage in the context of fair and equal benefit sharing. More generally, the web interface illustrated in Figure 4 enables the interactive exploration of DSI use in science by a features text search, data aggregation across the data warehouse and crosslinking to the original ENA records and ePMC records. It enables further complex filtering, grouping as well as visualition as charts, world map projects, and network diagrams. Based on the use cases provided in this CBD context, basic questions regarding DSI usage are visualized in different relationships to answer questions such as: Which countries use DSI? Which countries (groups) contributed DSI? Are there countries that use DSI but do not contribute DSI? This is implemented by four classes of use cases: *general overview of DSI, per country use of DSI, collaborative use in economic and hemisphere groups, world map projection, DSI citation network*.

**Figure 4.**
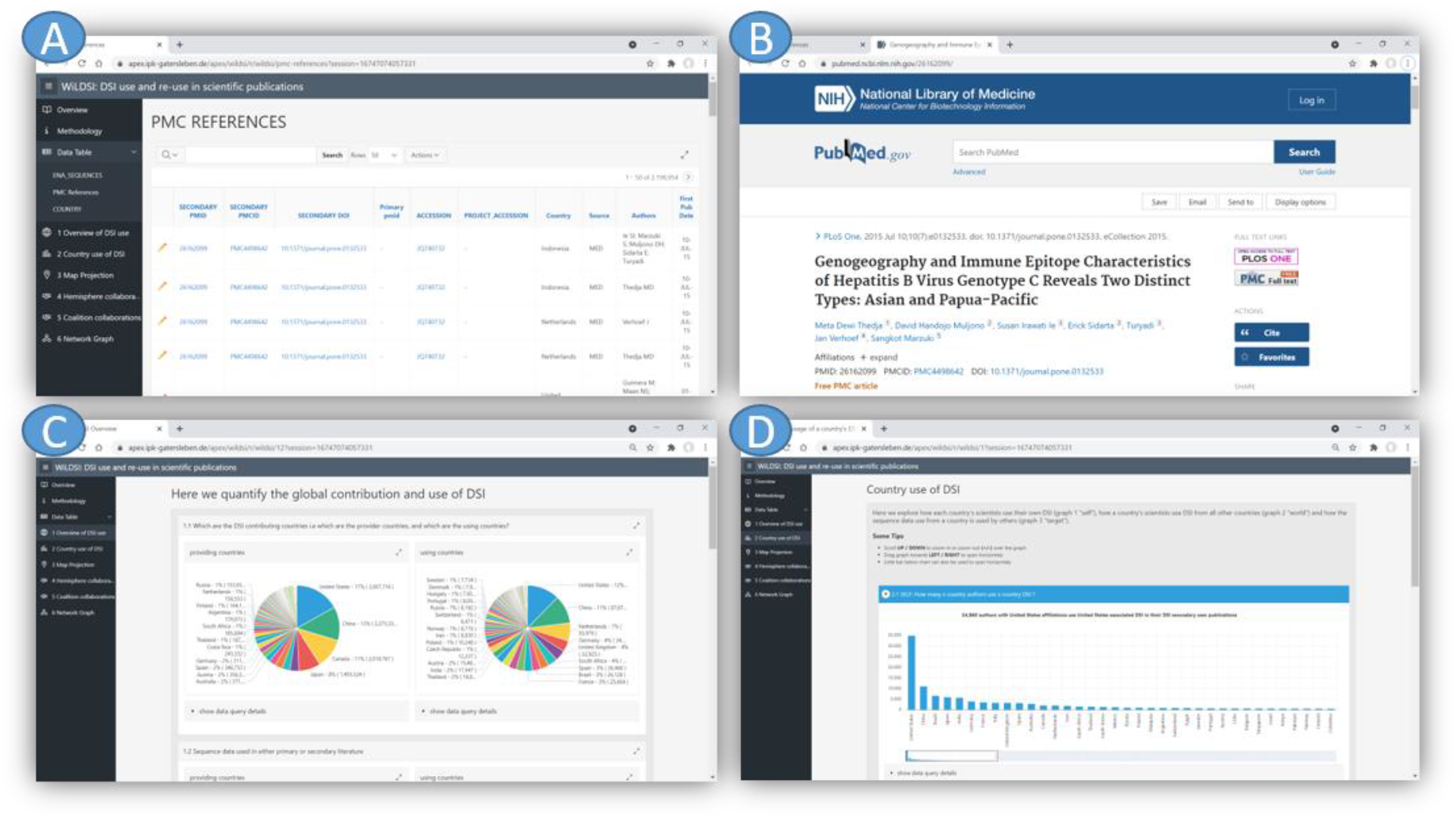
Screenshots of the WiLDSI Web Application. It consists of pages for (A) detailed data reports with integrated (B) drill down to original sources, (C) charts of DSI usage scenarios, (D) per country DSI use and contribution etc.

Another re-use scenario is to document the flow of DSI associated with genebank resources across the scientific value chain from seed storage to genetic analysis. A prominent example is the role of molecular passport data as an instance of DSI to characterize plant genetic resources (PGR). Genebank genomics is an emerging research field aiming at using high-throughput sequencing to characterize the genetic diversity in entire genebank collections [25]. Recently, marker profiles from reduced representation sequencing data were reported for more than 20,000 accessions of the German Genebank [26]. Whole-genome shotgun sequencing has been used to characterize the genome of 3,000 rice accessions at the International Rice Research Institute [27]. The approach provides a so-called molecular passport that enables tracking the identity of accessions, identifying redundancies and cross-link international genebanks [28]. For these reasons, molecular passport data is poised to become an essential component of working with PGR in research and breeding contexts.

Documenting the use of DSI associated with PGR would help genebank managers and administrators of genebank information systems monitor the use of their accessions in international research efforts and help justify the tremendous effort put into the maintenance and characterization of PGR in global genebanks. Documenting DSI could also help national authorities to enforce access and benefit-sharing schemes of the Nagoya protocol. The present enquiry into the status of DSI in public sequence archives has shown that sequence information of PGR is abundant, but tracing it back to the gene bank holdings it derives from, can be challenging. In the coming years, gene bank managers, genome researchers and bioinformaticians should develop and enshrine standards and protocols for linking DSI in archives such as EMBL-ENA to gene bank information systems and meta-databases such as EURISCO [29]. Work in this direction is underway in the EU-funded project AGENT [30].

DSI and their free accessibility are essential for all areas of the life sciences, including biodiversity research, food security, human health, biological conservation and many other disciplines or research areas. Some countries contributing DSI fear that direct access to the increasing amount of freely available sequence information may undermine benefit sharing schemes for genetic resources. A use of this data set supports evidence-based decision making in the context of international policy processes as well as global scale investigations into scientific use and re-use of NSD datasets and sub-disciplines thereof. Indeed, this article is intended as a companion paper for a timely publication on the policy implications of NSD (re-)use for DSI access and benefit-sharing discussions under the CBD in this issue.

For future studies, the examples above could be complemented by more detailed use cases including finer-grained groupings for data aggregation such as separation of genera, species and time ranges of publications. In combination with additional text classification techniques [32], it may be possible to cluster by research topics, e.g. considering only citations in paper involving, say, COVID-19 or plant pathogen resistance.

## Availability of source code and requirements

Project name: WiLDSI

Project home page: https://wildsi.ipk-gatersleben.de

Operating system(s): LINUX

Programming language: Oracle Application Express, Perl, Python3

Other requirements: HTML5 compatible web browser

License: GNU General Public License v3.0

All scripts used for data extraction are available from GitHub https://github.com/alakob/sequence-literature.

## Data Availability

The charts, maps, and data tables are available in an interactive web application at http://wildsi.ipk-gatersleben.de. The data tables are published as CSV files in the e!DAL-PGP repository [33] under the DOI 10.5447/ipk/2021/8. The SQL queries implementing the use cases are linked and documented alongside each chart within the web application.

## List of abbreviations

CBD: Convention on Biological Diversity
ITPGRFA: International Treaty for Plant Genetic Resources for Food and Agriculture
DOI: Document Object Identifier
EMBL: European Molecular Biology Laboratory
ENA: European Nucleotide Archive
ePMC: Europe PubMed Central
DSI: Digital Sequence Information - synonym for nucleotide sequence data in international policy circles
GR: Genetic Resources
INSDC: Nucleotide Sequence Database Collaboration
NSD: Nucleotide Sequence Data - synonym to DSI in a technical and database context
ORCID: Open Researcher and Contributor ID
PGR: Plant Genetic Resources
WiLDSI: German: “wissenschaftsbasierte Lösungsansätze für digitale Sequenzinformation”,
English translation: Science-based Approaches for Digital Sequence Information

## Ethics approval and consent to participate

Not applicable.

## Consent for publication

Not applicable.

## Authors’ contributions

Conceptualization: A.S., J.F., G.C., M.L.

Software: M.G., B.A., M.L., P.H., F. Z.

Data curation: U.H., J.F., M.L.

Investigation: A.S., J.F., U.H.

Supervision: M.L., A.S., J.F., G.C.

Writing original draft: M.G., M.L., M.M., A.S.

Writing review and editing: All authors

Funding acquisition: A.S., U.S.

## Acknowledgements

We thank H. Miehe, M. Oppermann and T. Münch for technical support and hosting the web application. We also thank the thousands of authors that generated the data and publications analyzed here especially to those committed to open access which enables this global overview.

## Footnotes

1 designated as secondary in CBD context

2 designated as primary in CBD context

3 “Ocean” is not equivalent to international waters under the United Nations Convention of the Law of the Seas (UNCLOS, where marine genetic resources and benefit-sharing are beeng discussed) but is in this context simply a consolidated term representing sampling in the marine environment.

4 Our queries used this URL pattern:https://app.dimensions.ai/discover/publication?search_text=AY924392&search_type=kws&search_field=full_search

5 Our queries used this URL pattern:https://www.lens.org/lens/scholar/search/results?q=AY924392&preview=true

